# Beneficial and Detrimental Effects of Nonspecific Binding During DNA Hybridization

**DOI:** 10.1101/2022.03.31.486645

**Authors:** Tam T. M. Phan, Tien M. Phan, Jeremy D. Schmit

## Abstract

DNA strands have to sample numerous states to find the alignment that maximizes Watson-Crick-Franklin base pairing. This process depends strongly on sequence, which affects the stability of the native duplex as well as the prevalence of non-native inter- and intra-molecular helices. We present a theory which describes DNA hybridization as a three stage process: diffusion, registry search, and zipping. We find that non-specific binding affects each of these stages in different ways. Mis-registered intermolecular binding in the registry search stage helps DNA strands sample different alignments and accelerates the hybridization rate. Non-native intramolecular structure affects all three stages by rendering portions of the molecule inert to intermolecular association, limiting mis-registered alignments to be sampled, and impeding the zipping process. Once in-register base-pairs are formed, the stability of the native structure is important to hold the molecules together long enough for non-native contacts to break.

## Introduction

The double stranded nature of DNA plays a crucial role in the storage, retrieval, and transfer of genetic information. These functions depend on the ability of DNA to separate into individual strands and then re-hybridize. The recognition of specific nucleotide sequences during hybridization also plays a fundamental role in medicine, biotechnology, and nanotechnology^1–4^ by enabling techniques like PCR^5–7^, DNA-based nanostructures (DNA origami)^8–12^, and methods for the diagnosis of diseases like HPV, HIV, and cancer^13–15^. The specificity in DNA hybridization is encoded in the complementary H-bonding patterns of the A-T and G-C Watson-Crick-Franklin (WCF) base pairs. However, the major driving force for duplex formation comes from the stacking of aromatic bases^16^. This contribution from base stacking has important consequences on the stability of double-stranded DNA helices. For example, there is a free energy penalty to initiate a helix due to the fact that base stacking cannot occur until consecutive base pairs have formed. Additionally, base stacking interactions depend on the identity of neighboring bases, resulting in helix stabilities that are strongly sequence dependent. Both the initiation and sequence dependent effects can be approximately captured using base pair interaction energies that account for nearest-neighbor bases ^5,17–22^.

While there are numerous tools to predict the thermodynamic stability of DNA duplexes^5,23–29^, the kinetics of hybridization are less well understood. This is a major limitation because kinetic limitations can prevent DNA molecules from finding the most stable state. The combinatorics of only having four bases means there is a 25% probability of an AT or GC match for each base pair forming in non-native alignments. Therefore, many random portions of DNA will be able to hybridize. While these states have a negligible effect on the equilibrium population of states^30^, they can prevent DNA from finding the correctly hybridized state within the required timescale and require special attention in the design of sequence-specific DNA oligomers^31^.

Experiments and theoretical analysis of long DNA molecules (kbp to Mbp) have shown that hybridization follows second order reaction kinetics that are limited by a slow nucleation step followed by a rapid zipping step^32^, and further analysis has explained how the hybridization rate scales with length, solvent viscosity, and sequence complexity^33^. Work on small molecules has provided high resolution views of hybridization. Simulations have revealed “inchworm” and “slithering” mechanisms in the search over intermolecular alignments^2,3,10,31,34–39^ and theory has explained the temperature dependence of the alignment search^40^. Single molecule experiments have shown that this search is very sensitive to sequence perturbations with a threshold of ∼ 7bp to initiate hybridization^41^. However, the sequences explored in these studies were too short to show the extensive non-native traps expected in longer molecules. Multi-state kinetics, indicative of traps and intermediate states, have been observed on sequences as short as 12 bp^42^ and non-Arrhenius temperature effects are suggestive of different hybridization mechanisms at high and low temperature^43^. Thus, there is a gap in our understanding of hybridization on scales between the short oligomer and genomic DNA. There is a considerable practical importance at the scale of tens to hundreds of base pairs because this range encompasses oligonucleotides used by PCR, CRISPR, DNA origami, and more. At this intermediate scale sequence dependent variations do not average out as they do on the genomic scale. Additionally, at this scale there is a significant probability of non-native base-pairing which increases the variability between sequences.

Here we present a theoretical model developed to explain the hybridization kinetics of 36 bp oligonucleotides studied by Zhang et al.^44^. Our model accounts for the competition between native and non-native base pairing during the random search over binding alignments with a three stage model^33^ using random walks to efficiently compute the effects of non-native interactions in each stage. We find that non-native *inter*molecular base pairs facilitate hybridization by increasing the lifetime of the encounter complexes in which the DNA strands search for native base pairs. In contrast, non-native *intra*molecular base pairs impede hybridization by creating barriers in the zipping process. However, intramolecular structure has a secondary beneficial effect of limiting the number of alignments that need to be searched.

## Methods

We introduce a model that accounts for three types of base pairing interactions.

### Native base pairs

Native base pairs, which we refer to as “in-register”, determine the stability of the final DNA duplex but only have a secondary influence on hybridization kinetics because most of the hybridization process involves searching non-native states.

### Mis-registered intermolecular base pairs

Misaligned DNA strands have a low probability of forming WCF matches and, therefore, have weaker binding affinity than in-register states. However, these misaligned states have a strong effect on hybridization kinetics due to the fact that most intermolecular collisions are out-of-register.

### Intramolecular base pairs

It is common for DNA strands to have self-complementary regions that allow the molecules to form intramolecular base pairs. While self-hybridization can lead to complex folds, especially in RNA, here we restrict our analysis to sequences that remain unstructured or only form simple “stem-loop” structures. We use the NUPACK software to predict these structures and compute their stability^23^ (Fig. 2). Such self-hybridized regions sequester a portion of the sticky bases, thereby increasing the probability that DNA molecules collide without hybridization. The intramolecular base pairs also must be broken to complete hybridization.

**Figure 1:**
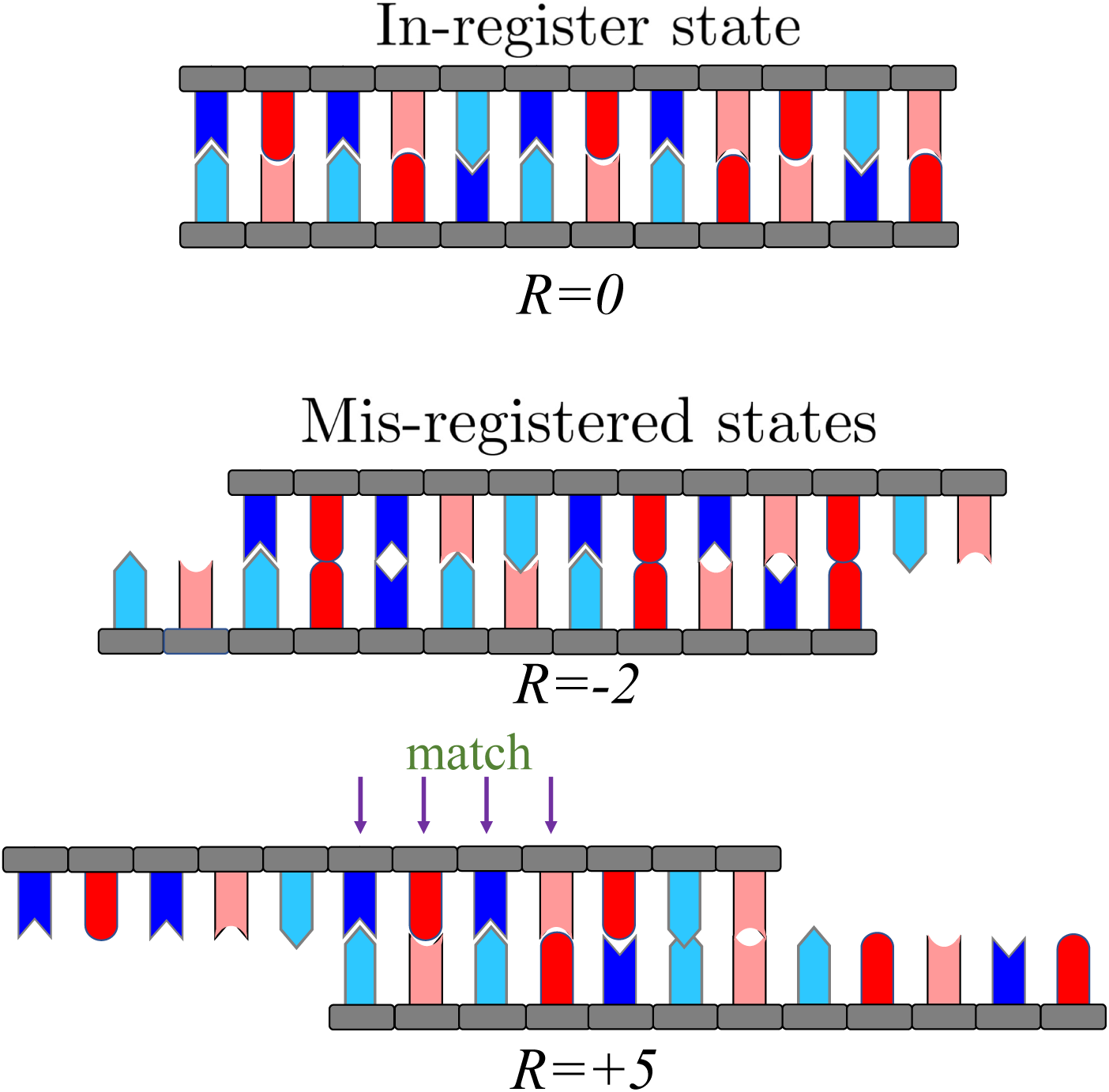
Cartoon representation of in-register and mis-registered states. In the in-register state, all base pairs follow the Watson-Crick-Franklin (WCF) rule in which A (dark blue) always pairs with T (light blue), and G (red) always joins C (light red). In contrast, in the mis-registered states, most base pairs are mismatches. However, because there are only four bases, many alignments will allow for the formation of a few WCF base pairs by random chance, resulting in a kinetic trap. The misaligned state at *R* = +5 shows a kinetic trap of four WCF base pairs.

**Figure 2:**
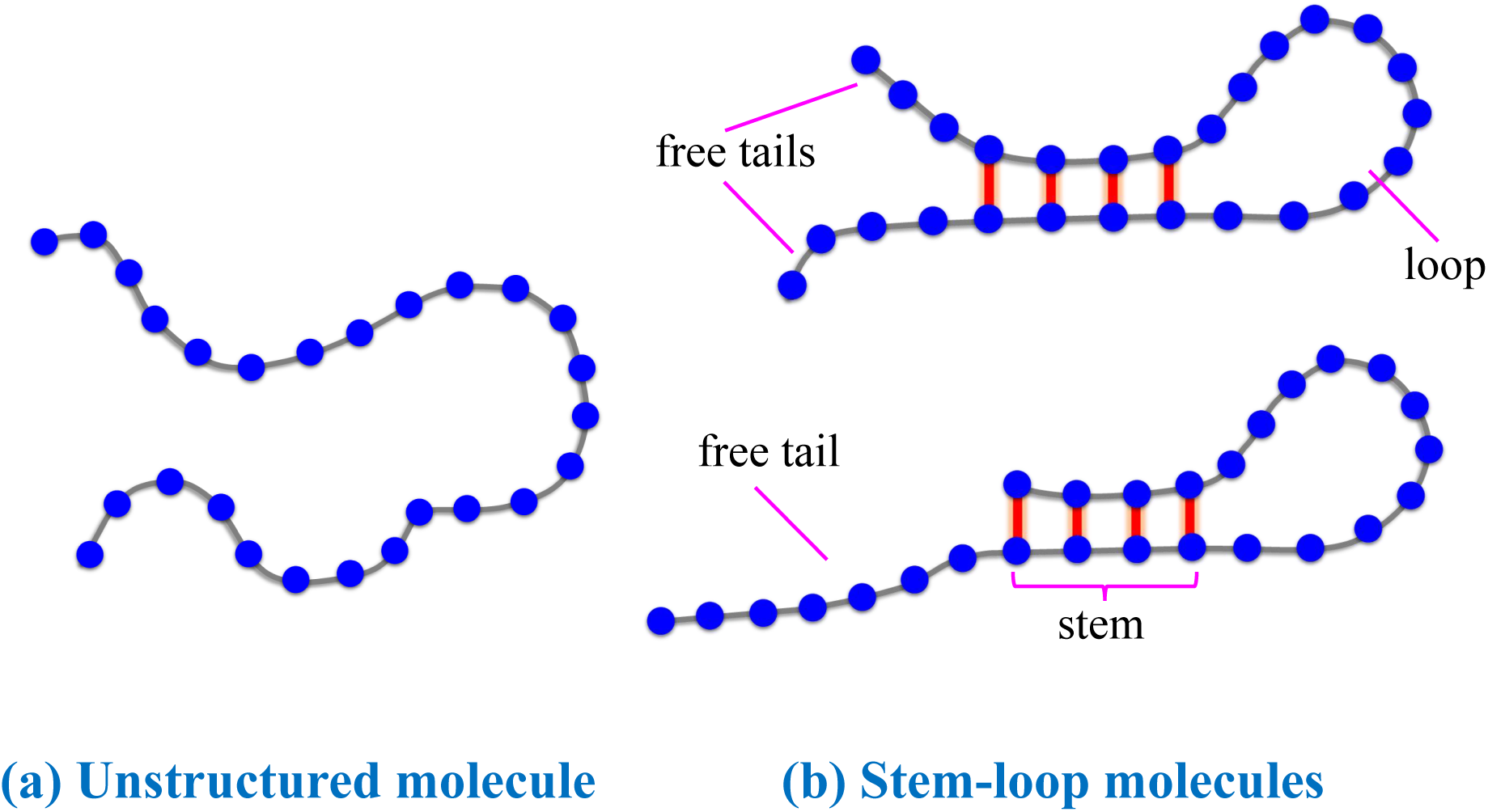
Cartoon representation of the structure of single DNA strands. (a) Unstructured molecules lack the self-complementarity required to form loop and stem regions. (b) Stem-loop structures occur when a sequence has a single self-complementary region. Intramolecular hybridization in this region results in the formation of a double stranded “stem” separating a single stranded loop and one or two free tails.

Figure 3 shows that metric specific to each of these kinds of interactions correlate poorly with hybridization kinetics when considered individually. The reason for this is that hybridization is a multi-stage process in which the bond types contribute differently to each stage.

**Figure 3:**
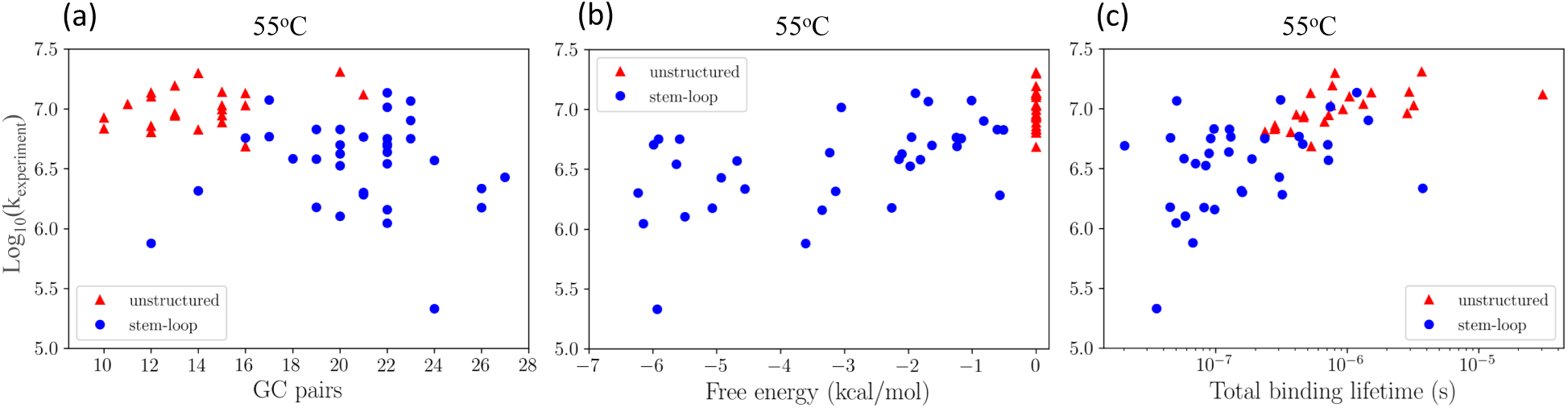
Hybridization kinetics correlate poorly with metrics that only consider one of the three types of base pairing. (a) Sequences with more GC content than AT content have a more stable hybridized state. However, this does not result in faster hybridization and, in fact, there is a weak trend suggesting that more GC content gives slower hybridization rates. (b) Sequences with more stable stem-loop structures (computed by NUPACK^23^) tend to hybridize more slowly than sequences with less stable stems or those that do not form stem-loop structures (red triangles). (c) The stability of misaligned states correlates with faster hybridization. The plot shows the sum of binding lifetimes in all misaligned registries. In all panels the scatter is larger than the trend (if present) indicating that multiple types of interactions must be accounted for to predict hybridization rates.

Our model includes these non-specific interactions using three stages, similar to those proposed by Niranjani et al.^33^. The stages, which we refer to as the diffusion stage, the registry stage, and the zipping stage (Fig. 4), each have a characteristic timescale as well as a probability that the system advances to the next stage.

**Figure 4:**
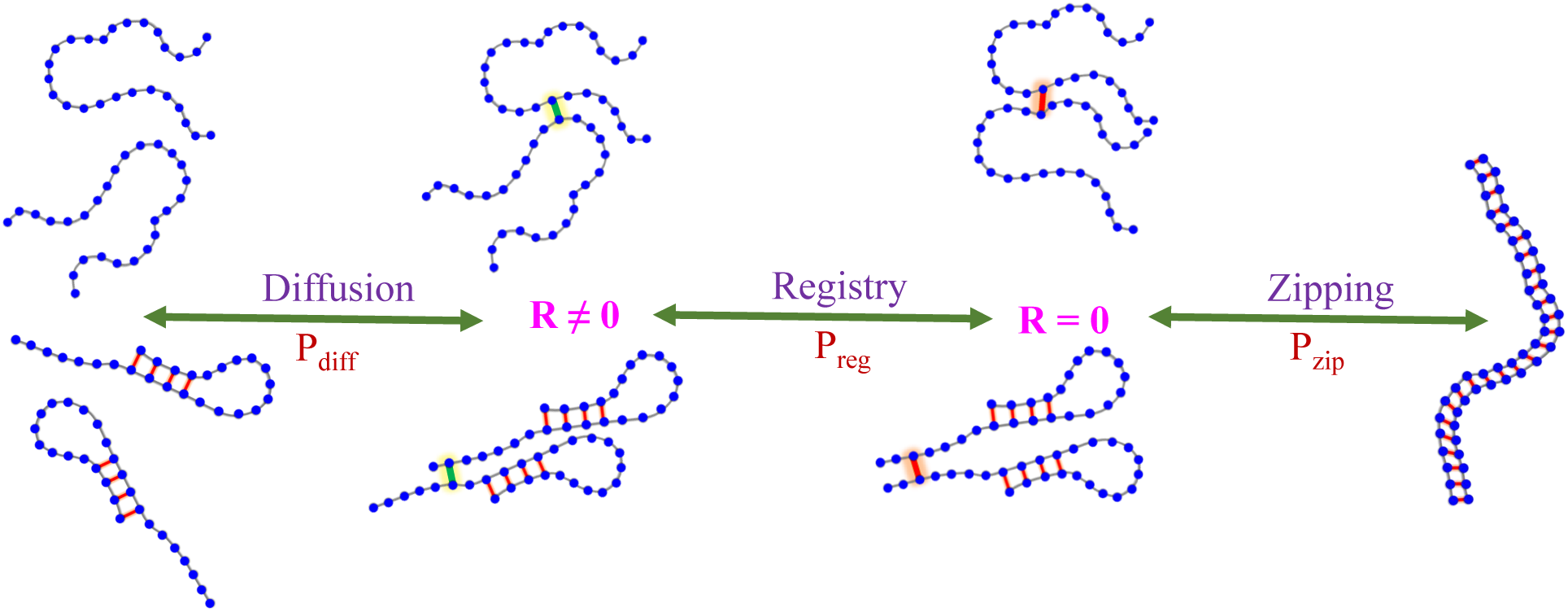
Diagram of the three stages of the DNA hybridization kinetics shown for both unstructured and single loop molecules. DNA strands go through the diffusion stage which usually results in an initial H-bond at *R* ≠ 0, the registry stage which results in the first *R* = 0 base pair, and the zipping state to obtain a fully hybridized state. In rare cases the DNA forms the initial bond at *R* = 0. In these events the registry stage is skipped and the DNA strands go to the zipping stage immediately after the diffusion stage.

### Diffusion stage

This stage describes molecules that have not formed any intermolecular base pairs. The characteristic time, *t*_diff_, is the inverse collision rate and *P*_diff_ is the probability that a collision results in the formation of an intermolecular base pair. We take *P*_diff_ = 0 in collisions where either of the colliding bases are involved in intramolecular stems. We also neglect the formation of intermolecular base pairs by bases in loop regions under the assumption that these bases are too constrained to properly stack.

### Registry stage

The registry stage is an encounter complex stabilized by non-native base pairs. We characterize the encounter complex using the registry variable *R*, which indicates how many bases a molecule needs to be shifted to form native base pairs (Fig. 1). The registry variable can take positive or negative values −*L*+1 ≤ *R* ≤ *L*−1, where *L* is the number of base pairs in a strand and *R* = 0 indicates the in-register (native) alignment^45,46^. Each registry has a binding lifetime *t*_reg_(*R*) in which the molecules search for in-register base pairs. *P*_reg_(*R*) describes the probability that the molecules held together by non-native base pairs in registry *R* form an in-register base pair before the non-native bonds dissociate.

### Zipping stage

Once two molecules form the first in-register base pair they advance to the zipping stage. *P*_zip_ indicates the probability that all native base pairs successfully hybridize before the molecules dissociate. This stage has an average duration *t*_zip+_ or *t*_zip−_ depending on whether the zipping stage ends with the formation of all in-register bonds or rupture of all in-register bonds before zipping completes, respectively. We assume that failure in either the registry or zipping stages results in the molecule returning to the diffusion stage.

#### Molecules randomly sample registries during hybridization

The hybridization rate is computed from the rate that molecules progress through these three stages. The first step is to write the time required for *N* intermolecular collisions.

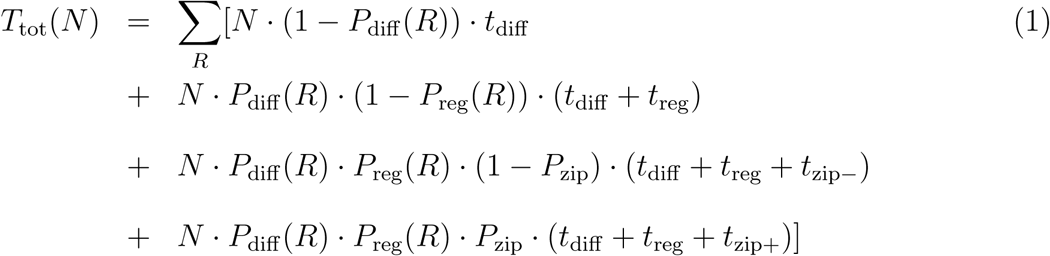

In Eq. 1 the first term represents failed collisions in which two strands are unable to bring bases into contact. *P*_diff_ (*R*) is the probability of forming a first bond in registry *R* after an inter-molecular collision. The second term accounts for molecules that form intermolecular bonds but do not form in-register bonds before dissolution. *P*_reg_(*R*) is the probability of forming a *R* = 0 base pair after forming at least one base pair in registry *R*. The third term describes events in which in-register bonds form but fail to reach the fully zipped state. *P*_zip_ is the probability for fully zipping after forming at least one *R* = 0 base pair. The last term describes successful collisions where two strands are able to arrive at the fully zipped state.

Noting that *T*_tot_(*N*) scales linearly with *N*, we can write *T*_tot_(*N*) = *NT*_tot_(1), where *T*_tot_(1) is the average time per collision. After *N* collisions there will be Σ_*R*_[*N* ·*P*_diff_ (*R*)·*P*_reg_(*R*)·*P*_zip_] successful events, so the hybridization rate is

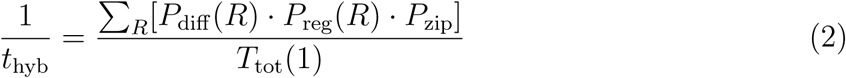

This expression can be simplified considerably in the low concentration limit where *t*_diff_ ⪢ *t*_reg_, *t*_zip_, which describes the 50 pM concentrations used in the experiments of Zhang et al.^44^. In this limit *t*_diff_ + *t*_reg_ + *t*_zip_ Δ *t*_diff_, which allows us to rewrite the hybridization rate as

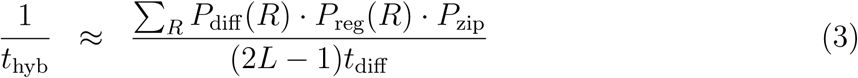

where (2*L* − 1) is the number of registries that need to be sampled.

#### The growth of the hybridized region is a random walk

To compute *t*_reg_, *t*_zip_, and *P*_zip_ we model the formation and breakage of base pairs as a first passage random walk^33^. To do this, we use the number of base pairs formed as the reaction coordinate ^47^. A system starting with *x* base pairs will evolve to *x* + 1 base pairs at a rate *k*_+_(*x*) or to *x* − 1 base pairs at a rate *k*_−_(*x*). Using these rates, we can write the probability of base pair formation as *p*_+_ = *k*_+_(*x*)/(*k*_+_(*x*)+*k*_−_(*x*)) and the probability of base pair breakage as *p*_−_ = *k*_−_(*x*)/(*k*_+_(*x*)+ *k*_−_(*x*)). The forward and backward rates are related by the detailed balance relationship

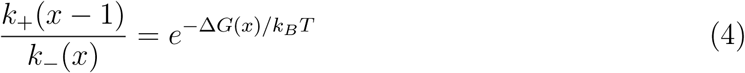

where Δ*G*(*x*) is the free energy change to form the *x*th base pair defined as Δ*G*(*x*) = *G*(*x*) − *G*(*x* − 1). We assume that bond formation *k*_+_ ≃ 10^9^*s*^−1^ is independent of sequence, while bond breakage is limited by the bond breakage energy *k*_−_ = *k*_+_*e*^Δ*G*^ where Δ*G* is the free energy of base pair formation computed with the nearest-neighbor free energies of Santa Lucia et al.^5,17–22^.

## Results and Discussion

### Diffusion stage

#### The duration of the diffusion stage is determined by random collisions

We estimate the diffusion time using the Smoluchowski formula for an absorbing sphere

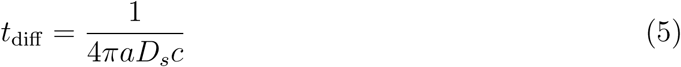

where *D*_*s*_ is the diffusion constant of the strands, *a* is the effective radius of the polymer coil, and *c* is the concentration of DNA. Inserting the Stokes-Einstein relation, *D*_*s*_ = *k*_*B*_*T/*6*πηa*, for the diffusion constant shows that the size dependence cancels when the target and incoming species are identical

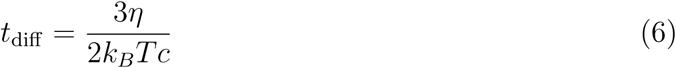

where *η* is the viscosity of the solvent.

#### The diffusion stage ends with the collision between two exposed bases

The sticking probability, *P*_diff_ (*R*), serves two purposes in our model. First, it provides a weighting factor to account for the fact that random collisions between DNA molecules are biased in favor of small |*R*|. To see this we observe that, of the *L*^2^ possible intermolecular base-base contacts, only (1,*L*) is consistent with registry *R* = *L* − 1. In contrast, there are *L* pairings: (1,1), (2,2),…(*L, L*) consistent with *R* = 0. Therefore, for unstructured molecules we have

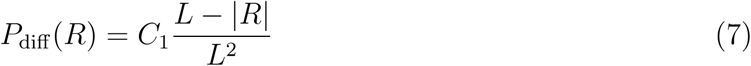

where *C*_1_ is a sequence-independent geometric factor that accounts for collisions in orientations incompatible with base pair formation (i.e., between phosphate backbones).

The second purpose of *P*_diff_ is to ensure that intermolecular base pairs only form between bases not previously engaged in intra-molecular base pairs. For stem-loop molecules this means the contact must be between free tails (we neglect binding in the loops). This limits registries that are possible because registries with |*R*| greater than the length of the tail cannot form base pairs. To handle this we need to apply Eq. 7 separately for each tail

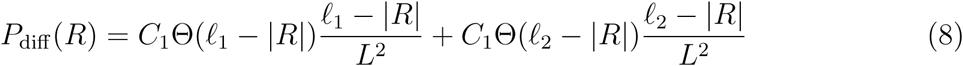

where *ℓ*_1_ and *ℓ*_2_ are the lengths of the two tails, we have neglected binding between opposite tails, *ℓ* − |*R*| is the number of possible collisions for each registry, and Θ is the Heaviside function defined as Θ(*n*) = 1 for *n >* 0 and Θ(*n*) = 0 for *n* ≤ 0. When a molecule with two free tails necessitates the use of Eq. 8, we also separately calculated *P*_reg_ and *P*_zip_ for each tail. In this case, Eq. 3 take the form

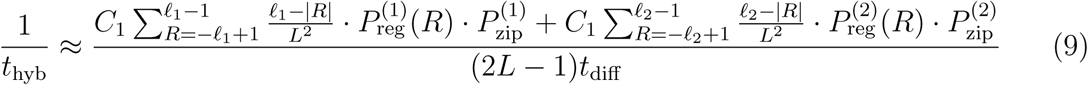

where the superscripts on *P*_reg_ and *P*_zip_ indicate the respective tail. The most important consequence of Eq. 8 is that molecules with extensive intra-molecular bonding have low values of *P*_diff_ (*R*) because the two strands can only form favorable interactions between the free tails.

### Registry stage

#### Non-native base pairs transiently hold two strands together

The registry stage is an encounter complex held together by non-native base pairs. The lifetime of the encounter complex is determined by the affinity of base pairs at the intermolecular contact. We compute the lifetime using Gillespie simulations^48^ in which the allowed moves are the formation and breakage of base pairs at either end of the intermolecular helix. These moves enter the simulations with rate constants given by Eq. 4 (see Appendix for details). The registry stage duration *t*_reg_ is computed as the average first passage time for a random walk starting at a single base pair to reach a state with zero base pairs.

Since the affinity of base pairs depends on both the sequence and alignment of molecules, there is a large variation in the registry stage lifetimes. Fig. 5 shows *t*_reg_ of an unstructured sequence as a function of the alignment *R*. Most alignments have very short binding lifetimes, less than 10 ns, due to the lack of contiguous complementary base pairs. However, misalignments of *R* = ±12 and *R* = ±17 allow for stretches of 7 and 6 WCF base pairs, respectively. As a result, these registries have binding lifetimes on the order of 1 *μ*s.

**Figure 5:**
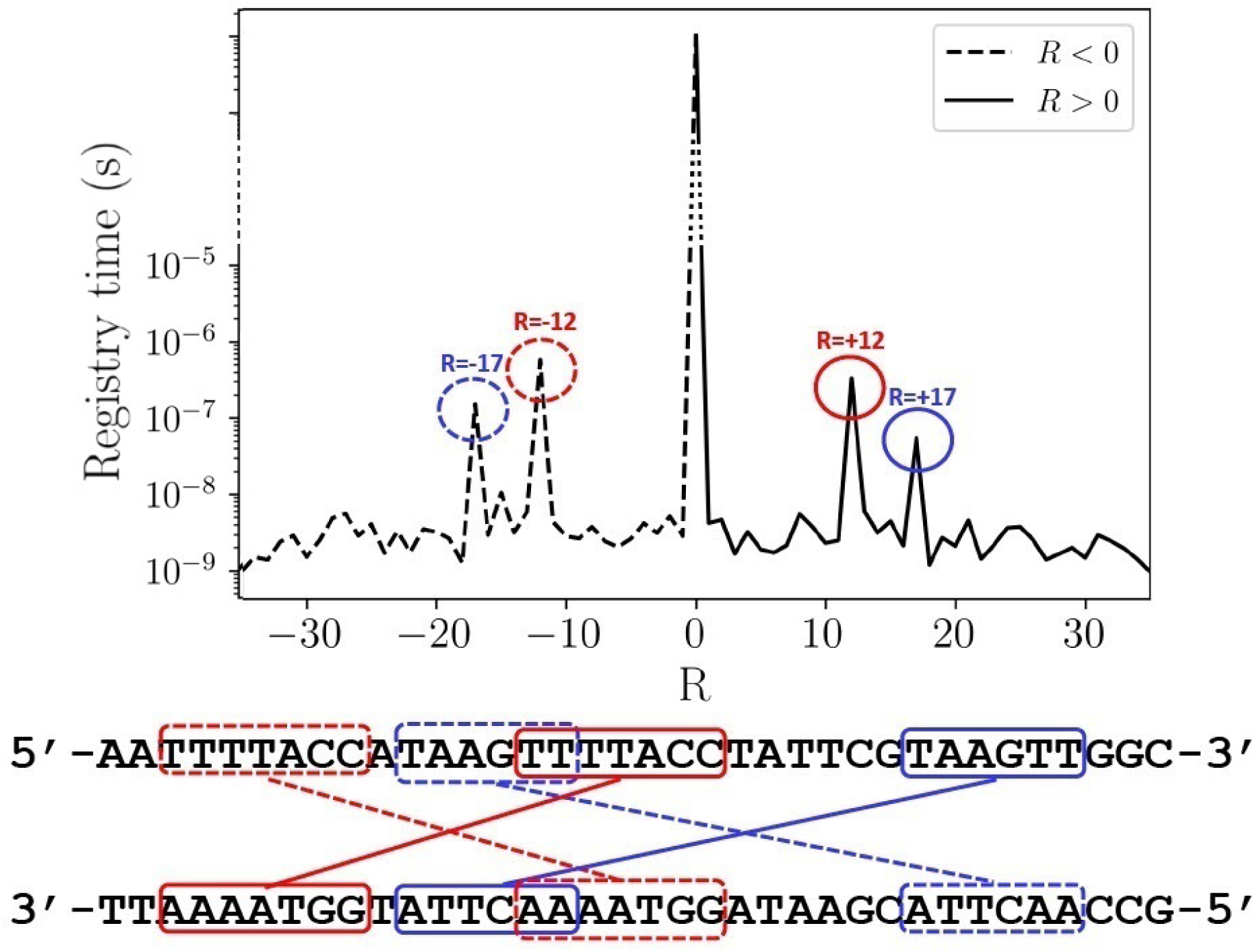
Comparison of the registry times of an unstructured sequence (S40) at 55°*C*. There are four peaks at registries *R* = ±17 (blue) and *R* = ±12 (red) which arise due to mis-registered WCF base pairs. These misaligned base pairs are indicated on the sequence below the plot with the same color codes (dashed lines indicate *R* < 0). The long lifetime of these states increases the probability that the molecules find in-register base pairs.

**Figure 6:**
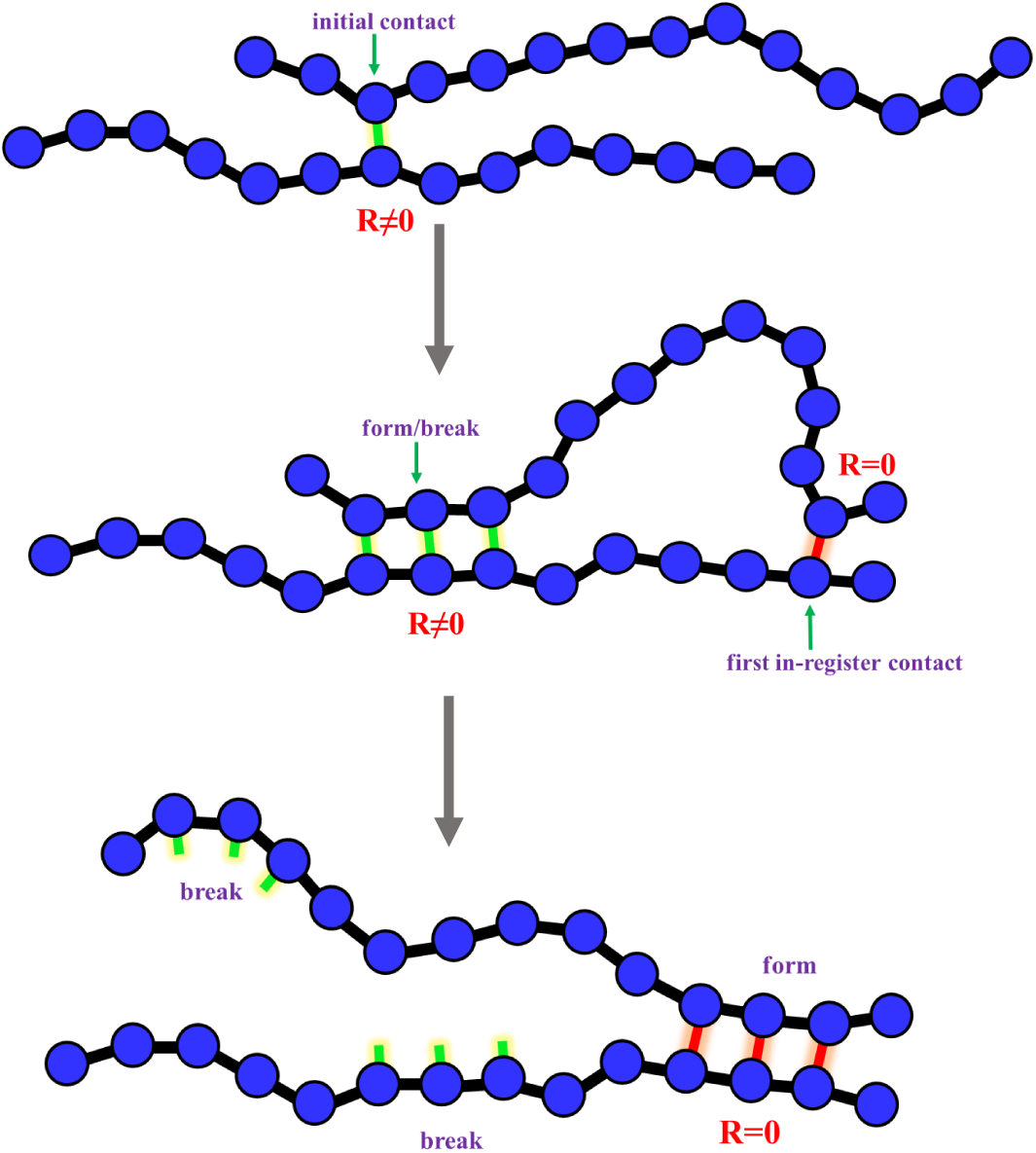
Cartoon representation of alignment searches of DNA strands. After forming the initial contact at *R* ≠ 0, the DNA strands may form or break bonds around the initial contact. In the meantime, the free regions around the initial contact fluctuate to search for *R* = 0 positions and have the first in-register contact. They may form or break WCF bonds around that in-register bond, but formation is more favorable. In contrast, misaligned bonds are less stable and have a relatively short lifetime.

#### DNA strands search for in-register base pairs during the registry search

While the molecules are held together by mis-registered base pairs, the free tails are free to fluctuate and search for in-register contacts via “inchworm” moves. We expect that the tails will come into contact on a time scale comparable to the Zimm time, *τ*_*Zimm*_, which describes the relaxation modes of a polymer in dilute solution^49,50^. Each tail-to-tail contact provides an opportunity for the molecule to find in-register base pairs. The probability of success depends on both the amount of time the molecules are held together, which determines the number of attempts, and the length of the free tails, which determines the probability a given attempt is successful. The number of attempts during the registry stage is *t*_reg_(*R*)*/τ*_*Zimm*_. To estimate the probability that one of these attempts is successful, consider two unhybridized segments of length *ℓ* joined by a single base pair in registry *R*. This leaves *ℓ* −1 bases on each strand available to form new intermolecular contacts (we neglect fluctuations in the number of non-native base pairs), so the number of possible new contacts is (*ℓ* − 1)^2^. However, only *ℓ* − 1 of these contacts are in-register. Furthermore |*R*| of the in-register contacts involve one of the strands folding back across the original base pair. After such a fold, any new base pairs will have parallel backbones (e.g., 5’ → 3’ with 5’ → 3’), which is incompatible with hybridization. Therefore, the probability that a random inter-strand contact is compatible with in-register hybridization is

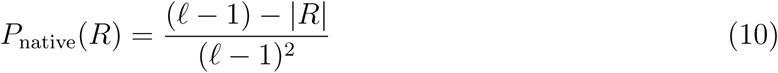

where (*ℓ* − 1) − |*R*| is the number of possible in-register base pairs and (*ℓ* − 1)^2^ is the total possible base pairs the molecules can search.

If *t*_reg_(*R*)*/τ*_*Zimm*_ is small *P*_reg_(*R*) can be approximated by *t*_reg_(*R*)*P*_native_*/τ*_Zimm_, however, if *t*_reg_(*R*) is large this expression can exceed unity. This is because the small time approximation allows multiple successful attempts in the allotted time. The desired quantity, *P*_reg_(*R*), is the probability that at least one tail-to-tail contact finds the in-register state during *t*_reg_. This is equivalent to *P*_reg_(*R*) = 1 − *P*_fail_(*R*), where *P*_fail_(*R*) is the probability that all attempts fail. The probability of a single attempt fail is 1 − *P*_native_(*R*), so

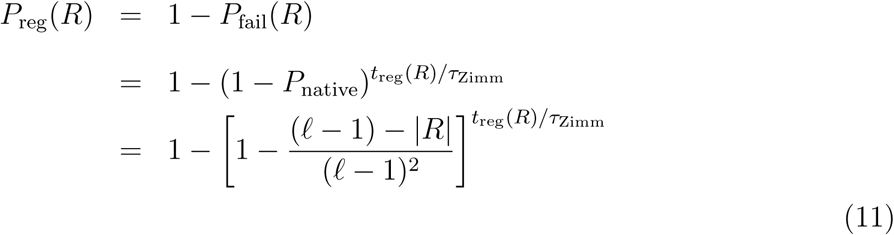

where *ℓ* = *L* for unstructured molecules and the length of the relevant tail, *ℓ*_1_ or *ℓ*_2_, for stem-loop sequences.

The Zimm time is given by *τ*_Zimm_ ∝ *ηR*^3^*/kT* ^49^, which can be rewritten as

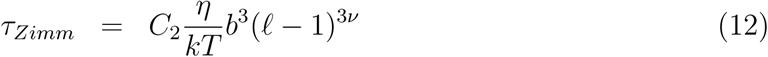

where *C*_2_ is a constant and *R* = *bn*^*ν*^ is the Flory radius with *b* ≈ 0.3 nm and *ν* = 0.588.

The rare cases where the initial intermolecular contact forms an in-register base pair can be described by *P*_reg_(0) = 1 and *t*_reg_ = 0, which means the molecules proceed directly from the diffusion stage to the zipping stage, skipping the registry stage.

Fig. 7 compares the computed *P*_reg_ and *t*_reg_, averaged over registries, for all sequences in the data set. We note two trends. First, longer binding lifetimes correlate with a higher probability of finding the in-register state. This is due to the fact that long-lived encounter complexes have more opportunities for the tails to collide and explore potential registries. Second, when the binding lifetimes are equivalent, the probability of finding in-register bonds is greater when part of the molecule is self-hybridized (orange squares and blue circles). This is because the intramolecular helix restricts the number of mis-registered base pairs that need to be sampled. In fact, unstructured regions shorter than 11 bases (orange squares) have > 70% chance of finding the in-register state with binding times of just a few nanoseconds. However, unstructured sequences (red triangles), which must search 36^2^ contacts, have less than 50% chance even when the binding lifetimes approach a microsecond.

**Figure 7:**
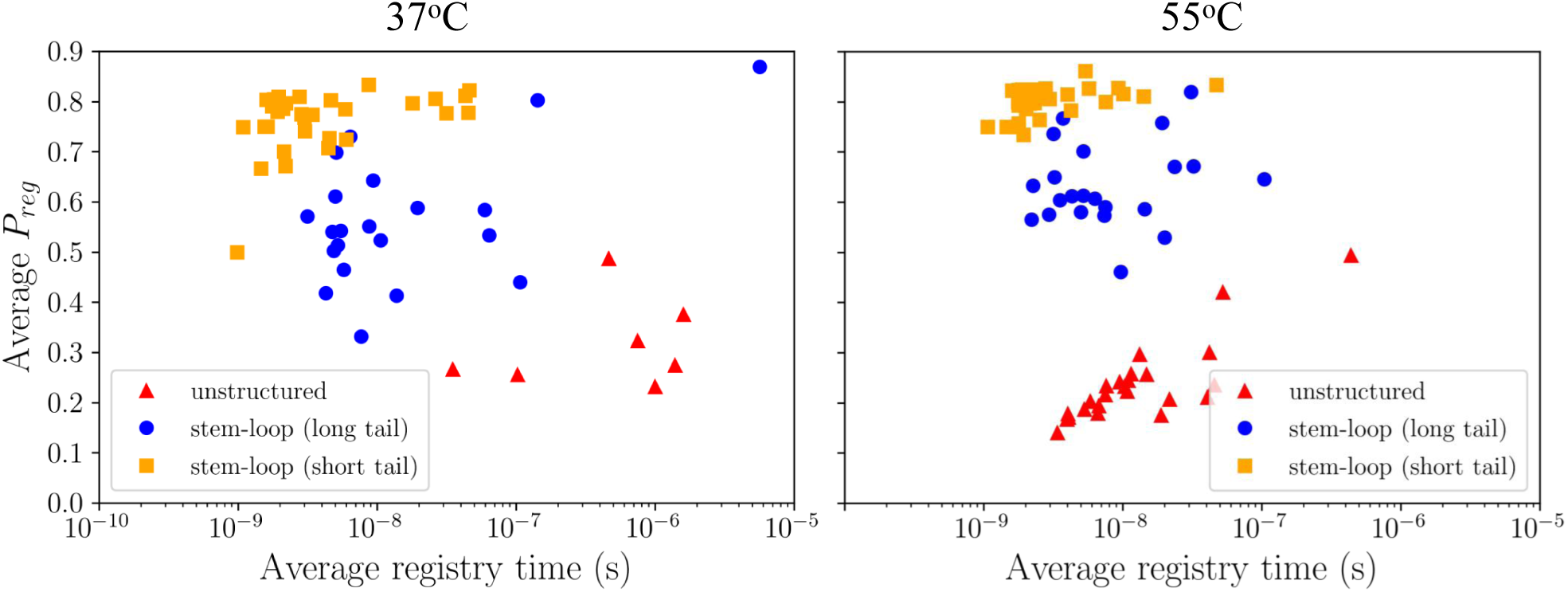
The calculated probability of finding the in-register state, *P*_reg_, averaged over registries with nonzero binding probability (see Eq. 8). Unstructured molecules (red triangles) have lower probability of finding the in-register state than stem-loop molecules (blue circles, orange squares) due to the greater number of intermolecular contacts that need to be searched. Similarly, tails with 12 or more bases (blue circles) have a lower probability than tails with 11 or fewer bases (orange squares).

### Zipping stage

#### The first in-register bonds provide an anchor to hold the strands together while hybridization completes

After forming the first in-register bonds, the DNA strands enter the zipping stage. The dynamics of the zipping stage are similar to the registry stage in that both phases involve the formation and breakage of base pairs at the edge of a growing helix. The success of this stage is determined by whether the native base pairs are able to form across the full length of the DNA before the molecules detach. There are two obstacles to a successful outcome. The first is that the initial in-register base pairs provide very little stability, so the nascent helix is prone to dissociate before the “toehold” grows large enough to form multiple base stacking interactions. Second, if the molecules have intramolecular base pairing, the stems will need to be broken before native base pairs can form in these regions. These obstacles depend only minimally on the specific sequence of the molecule, which allows us to we obtain an analytic form for *P*_zip_ that captures the most important characteristics of the zipping stage.

To solve for *P*_zip_ we introduce the quantity *P*_zip_(*x*), which is the probability that a pair of molecules with *x* base pairs forms the last (*L*th) base pair before falling apart. The quantity *P*_zip_ in Eq. 1, which describes the probability of the full zipping after forming the first base pair is given by *P*_zip_(1). *P*_zip_(*x*) satisfies the recursion relation

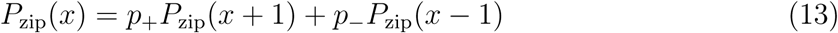

which says that a random walk starting with *x* base pairs will proceed to *x*+1 with probability *p*_+_ and to *x* − 1 with probability *p*_−_ ^51^. In the continuum limit Eq. 13 takes the form of a one-dimensional convection-diffusion equation

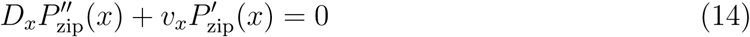

where the drift velocity and diffusion coefficient are given by

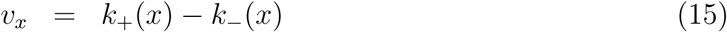

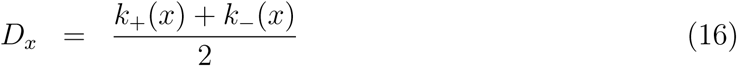

If *x* describes a region of the molecule without intramolecular bonding, then the new base pairs are favorable (Δ*G*(*x*) < 0) so Eqs. 4 and 15 give *v >* 0. As a result, there is a strong bias for the helix to grow. However, in regions with intramolecular helices, two non-native bonds must be broken for each intermolecular bond that is formed. Since the two intramolecular bonds that are broken have the same sequence as the single intermolecular bond, they incur an energetic cost −2Δ*G*(*x*). Therefore, the net change is Δ*G*(*x*) − 2Δ*G*(*x*) = −Δ*G*(*x*) *>* 0 indicating that the formation of intermolecular bonds is unfavorable in the stem region. As a result, zipping within regions of intramolecular structure has a bias in the negative direction (*v* < 0). Importantly, sequence-dependent energetics of hybridization in stem regions are equal in magnitude and opposite in sign to that of unstructured regions. These energetic regimes are accounted for by taking *P*_zip_(*x*) to be a piecewise continuous function with the boundary conditions shown in Fig. 8.

**Figure 8:**
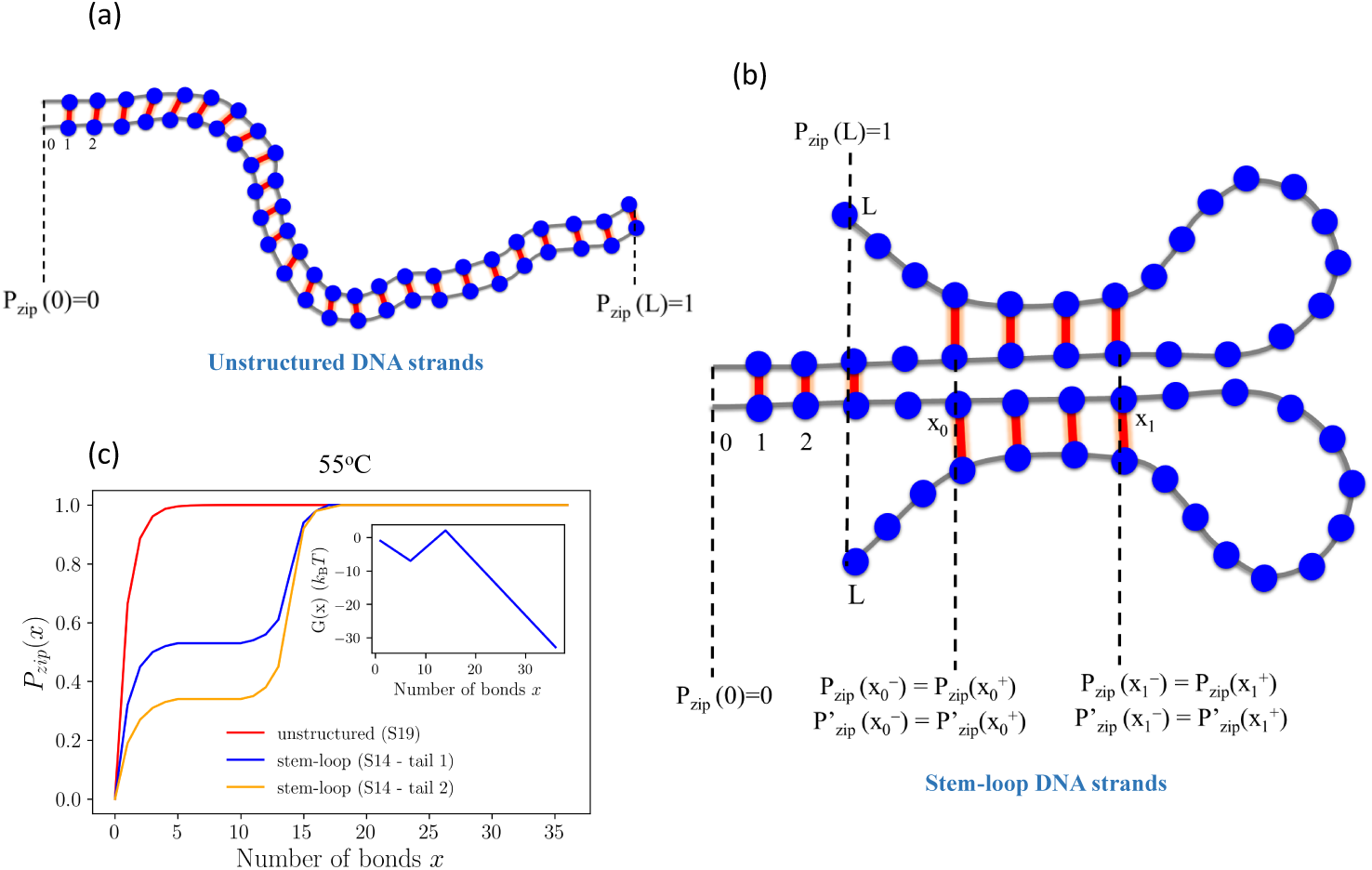
Cartoon representation of the boundary conditions for unstructured and stem-loop sequences in solving for *P*_zip_ (Eq. 14). (a) Unstructured molecules have two boundary conditions indicating the failure of zipping *P*_zip_(0) = 0 and successful zipping *P*_zip_(*L*) = 1. Stem-loop structures have the same boundary conditions at *x* = 0 and *x* = *L*, but have additional internal boundaries to ensure the function *P*_zip_(*x*) is continuous across the three regions. The first region 0 < *x* < *x*_0_, the “toehold”, describes base pairs within the free tail region. The second region *x*_0_ < *x* < *x*_1_ is where intramolecular stems are replaced with intermolecular base pairs. The third region begins when the last intramolecular base pairs are broken and encompasses *x*_1_ < *x* < *L*. In this region the formation of intermolecular base pairs is again favorable and *v >* 0. (c) The zipping probability is plotted as a function of *x* for an unstructured and a stem-loop sequence. *P*_zip_(*x*) rapidly approaches unity for the unstructured sequence reflecting the fact that dissociation is unlikely after ∼3 base pairs have formed. The stem-loop sequence has two tails of length *ℓ*_1_ = *ℓ*_2_ = 7 and a stem region that also has length 7. The zipping probability has a flat region for 3 < *x* < 12 because the drift velocities on both sides of this region tend to push the random walk toward a local minimum in the free energy at *x* = 7. The free energy profile for tail 1 is shown in the inset. The difference in *P*_zip_(*x*) between the two tails of S14 come from the greater stability of tail 1 (see Table 1 and accompanying text).

Figure 8c shows the solution to Eq. 14 for an unstructured sequence and both tails of a stem-loop sequence. The local diffusion coefficient and drift velocity are obtained by averaging the nearest-neighbor binding energies within each of the three regions (or entire sequence for the unstructured molecule). The zipping probability grows rapidly with the number of formed base pairs, *x* for the unstructured sequence reflecting the fact that failed zipping events are dominated by cases where the molecules have yet to establish a stable toehold. In contrast, the stem-loop structure shows an intermediate plateau over the region encompassing the anchoring tail and the intramolecular stems. This feature is a result of a local energy minimum where the intermolecular helix meets the bases of the intramolecular stems (Fig. 8c inset). Deviations away from this minimum require either the rupture of intermolecular base pairs or intramolecular stems, both of which are unfavorable. Therefore, the local drift velocities will tend to return random walks with 0 < *x* < *x*_1_ to the minimum at *x* = *x*_0_. This tendency to return to the base of the stems explains why the zipping probability is flat between *x* = 1 and *x*_1_. However, the probability jumps dramatically when the stems fully dissolve because there is no further impediment to zipping.

**Table 1:**
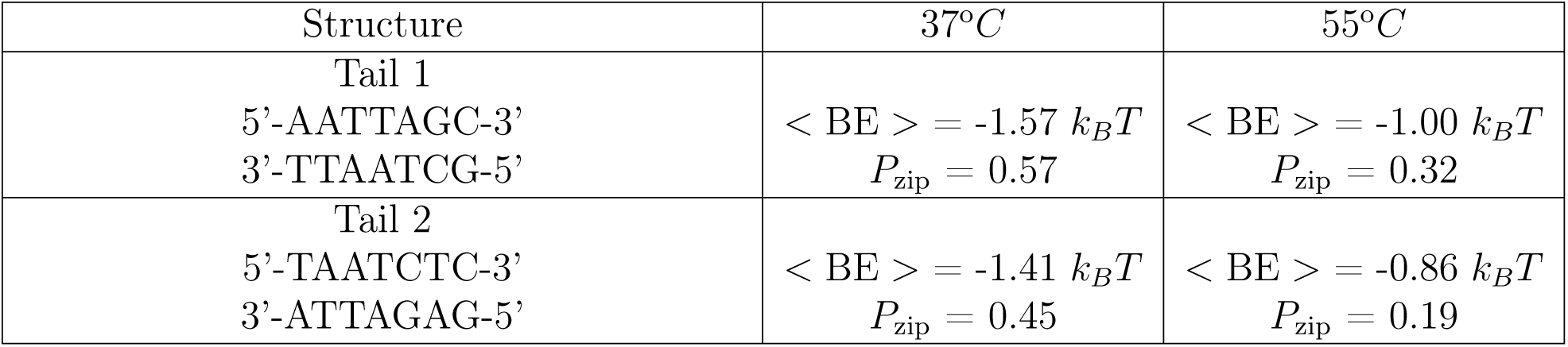
Average binding energy per base pair and *P*_zip_ of the two free tails of S14. Despite identical AT/GC composition, tail 1 has a stronger binding energy due to more favorable nearest-neighbor interactions. This gives tail 1 a longer binding lifetime and a higher probability of completing the zipping process.

#### Successful zipping is a competition between the stability of the anchoring base pairs and the structure that must be broken

The most important factor influencing *P*_zip_ is the stability of the intramolecular structure that must be broken compared to the stability of the intermolecular helix that holds the molecules together ^52,53^. Fig. 9 plots *P*_zip_ versus the ratio of the length of the unstructured tail to the length of the intramolecular stems. The figures shows that *P*_zip_ transitions from large values ∼ 70% when the intramolecular stems are shorter than the toehold intermolecular bonds, to small values (< 10%) when the stems are longer than the toehold. This observation suggests that *P*_zip_ is a competition between the lifetime of the intermolecular toehold and the lifetime of intramolecular stems. This interpretation is supported by the interesting case of sequence S14, which has two free tails of length *ℓ*_1_ = *ℓ*_2_ = 7 nucleotides. Furthermore, each tail has five A/T nucleotides and two G/C nucleotides. However, *P*_zip_ for tail 1 is greater than *P*_zip_ for tail 2 because the nearest-neighbor interactions in tail 1 leads to more favorable binding energy (Table 1, Fig. 8c). Therefore, tail 1 has a longer binding lifetime giving it a better chance for hybridization to progress through the intramolecular stems.

**Figure 9:**
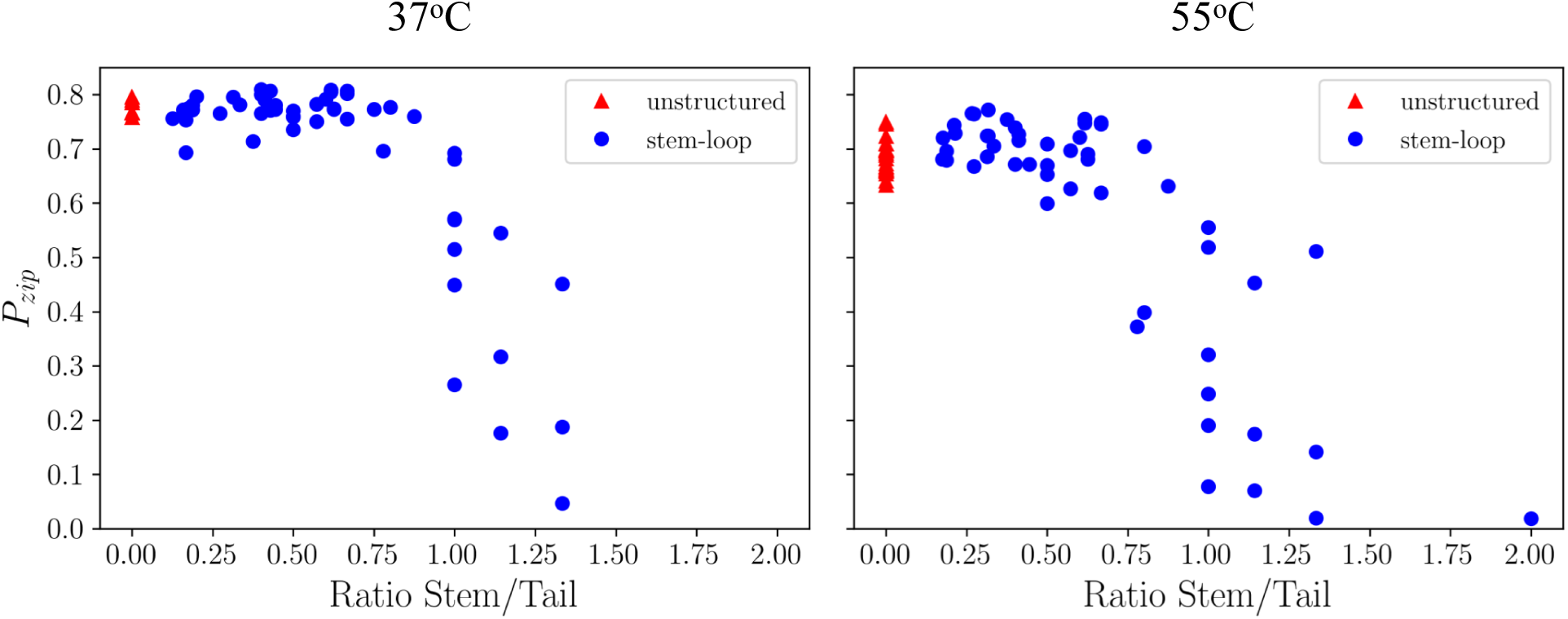
The zipping probability computed from the theoretical model plotted against the ratio of the number bases in the intramolecular stem to the number of bases in the free tail. The red triangles represent unstructured sequences, while the blue circles indicate stem-loop sequences. The probability drops dramatically when the intramolecular stems are longer than the intermolecular helix holding the molecules together suggesting that *P*_zip_ is a competition between the lifetime of these two structures.

#### In-register molecules zip rapidly unless there is intramolecular structure

The zipping time can be subdivided into events for which the zipping is either successful (*t*_zip+_) or not (*t*_zip−_). These times were computed via Gillespie simulation, as described above for the registry time, and plotted in Fig. 10. The failure times are generally shorter than the successful events due to the fact that many failure events occur shortly after initial binding. Fig. 10 also shows that longer intramolecular stems slow the zipping time because these system require an energy excitation large enough to rupture the stems.

**Figure 10:**
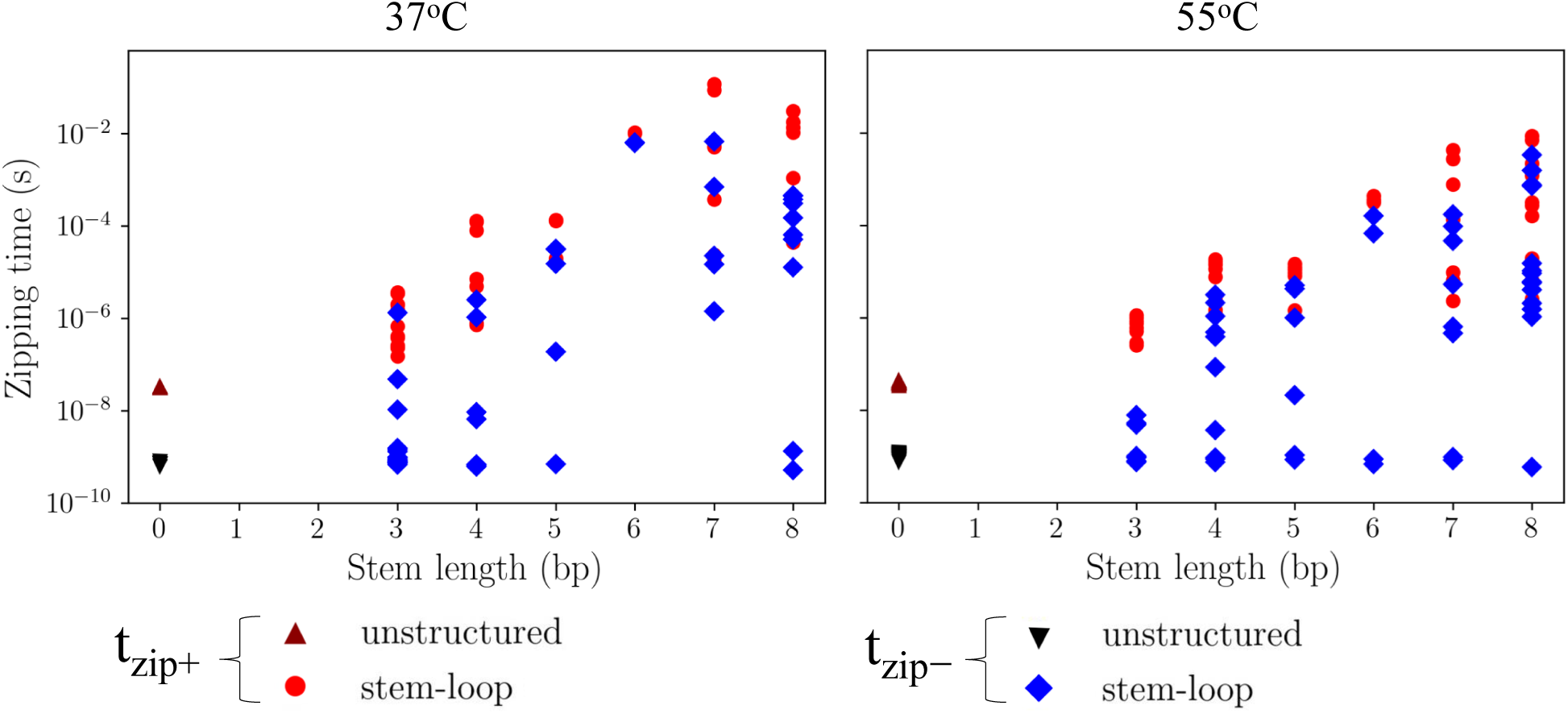
Calculated times for successful (*t*_zip_+) and failed (*t*_zip_ −) zipping stages plotted as a function of intramolecular stem length. Longer stems correlate with longer zipping times because it takes longer to get the energy fluctuation needed to melt the stem. The failure times show a weaker correlation because this outcome is determined primarily by the length of the unstructured tails.

### Diffusion is slower than the registry search or zipping

Figure 11 compares *t*_diff_, *t*_reg_, and *t*_zip_ for two molecules at 55°*C* with opposite characteristics. Sequence 12, has the longest zipping time in our calculations due to a very stable stem-loop structure. However, the calculated *t*_zip_ ≃ 10^−2^ s is still about 3 orders of magnitude faster than the time between diffusive collisions at the experimental concentration of 50 pM. This plot predicts that the hybridization rate will increase with concentration up to approximately 10^−8^ M, at which point the zipping time will become rate limiting. Sequence 73 shows the opposite extreme. Since S73 lacks intramolecular structure, zipping is unimpeded and completes in less than 10^−7^ s. In addition, S73 has a very long registry stage due to mis-registered states with lifetimes on the order 10^−7^ s. In this case the hybridization rate will not plateau until concentrations on the order of 10^−2^ M when mis-registered traps become limiting. To capture the plateau of hybridization rates at high concentrations, it is necessary to use Eq. 2 and not the approximate form Eq. 3.

**Figure 11:**
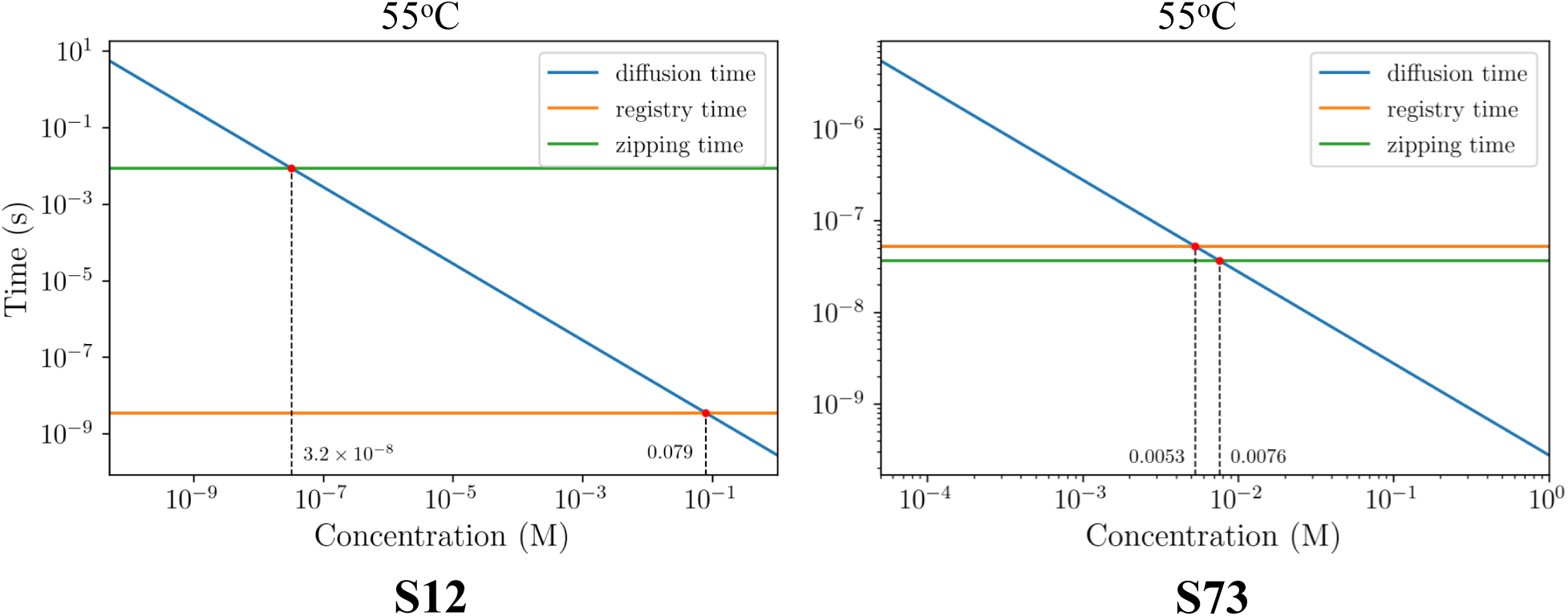
Comparison between the diffusion time (blue line), the average registry time (orange line), and the zipping time (green line) as a function of DNA concentration at 55°*C*. Sequence 12 (left panel) has a very stable stem-loop structure and, consequently, has the longest zipping time in our simulations. The unstructured sequence 73 (right panel) represents the sequence having the shortest zipping time. The red dots show the concentrations at which the diffusion time equals the registry time and the zipping time. These concentrations are much larger than the 50 pM concentration used in the experiments of Zhang et al., which justifies the use of the approximate rate expression Eq. 3.

### The wide range of hybridization rates comes from both native and non-native base pairing

The hybridization rate computed with our model is compared to the experiments of Zhang et al.^44^ in Fig. 12. There is a strong correlation between the computed and experimental values over nearly three orders of magnitude. Importantly, our model captures temperature dependent effects as well as the presence or absence of intramolecular structure with the same set of parameters.

**Figure 12:**
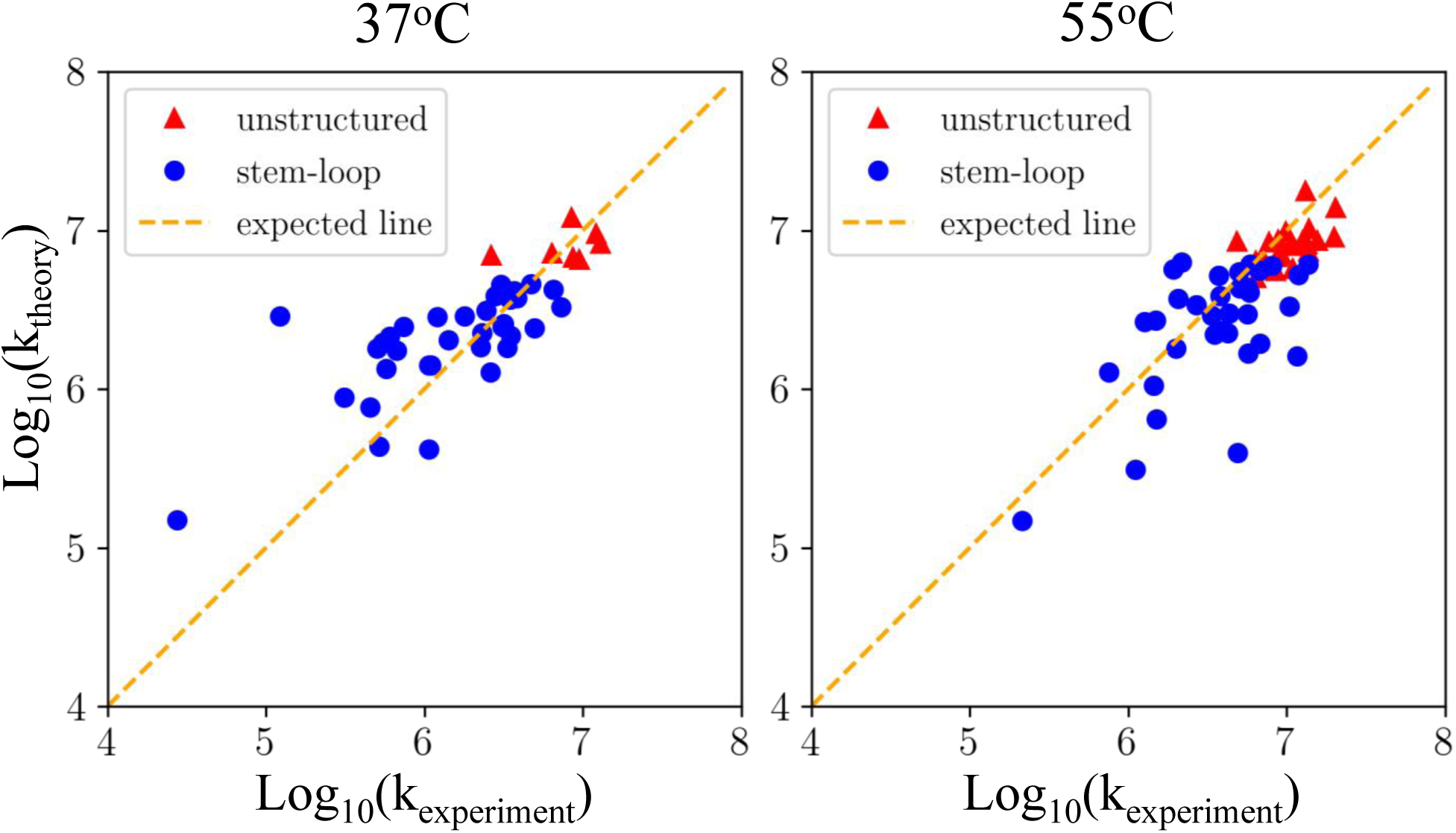
Comparison between the theoretical model and experimentally measured DNA hybridization rate at 37°*C* and 55°*C*. The hybridization rates *k*_*experiment*_ and *k*_*theory*_ are in units of *M* ^−1^*s*^−1^. The red triangles represent unstructured sequences, the blue circles indicate stem-loop sequences, and the dashed line shows optimal agreement between theory and experiment. In both cases, the theory fits well with the experiment. The free parameters, determined by a least squares regression, are the sequence independent geometric factor in Eq. 8 (*C*_1_ = 0.7), and the prefactor in the expression for the Zimm time Eq. 12 (*C*_2_ = 0.2).

Based on these results, we can understand the effects of both native and non-native base pairing on the hybridization rate. Typically in biomolecule self-assembly, native bonds have negligible contribution to the assembly kinetics because most of the time is spent searching the non-native ensemble^47,54–58^. However, here we find that native bonds play an important role in holding molecules together long enough for the hybridization to progress through non-native stem-loop structures. Another interesting finding is that non-native intermolecular bonds are beneficial for hybridization because they stabilize encounter complexes that facilitate the search for native contacts. This differs from the alignment search during amyloid protein aggregation where non-specific interactions seem to uniformly inhibit assembly kinetics^45,47,59^. We speculate that the difference is due to the strong specificity of the WCF base pairing, which limits non-native bonding to small regions with complementary sequences. This limitation on non-specific bonding frees the rest of the molecule to search for native contacts. In contrast, the backbone H-bonds stabilizing amyloid structures are more promiscuous, which allows the non-native bonding to propagate along the molecule and limits the length of free tails that can perform “inchworm” moves. Finally, we find that non-native intramolecular base pairs have both a beneficial effect in restricting the registries to be searched, and an inhibitory effect in creating a barrier to complete hybridization. The inhibitory effect is only significant, however, when the intramolecular stem is longer than the neighboring regions that hold the molecules together. Therefore, short intramolecular stems are likely to be mostly beneficial. The varied effects of these three types of bonding explain why hybridization rates correlate poorly with metrics like native stability, intramolecular structure, or the lifetime of mis-registered states (Fig. 3).

There are several approximations in our model that may contribute to the scatter in Fig. 12. These include our neglect of fluctuations in the size of stem structures or multiple inchworm moves in a single hybridization attempt. Another approximation that is likely to be significant is our neglect of intermolecular interactions between the loops formed by intramolecular stems. Clearly, the potential for WCF base pairing by these loops will strongly depend on the size of the loop and further investigation will be necessary to identify how long the loop needs to be before it contributes significantly to hybridization. In addition, there are other factor that come into play near surfaces^60–64^ or *in vivo*^65^.

It is interesting that our model performs as well as it does without accounting for the sliding of two unhybridized molecules past each other^2,3,10,31,34–39^. Intuitively, one might expect that such “slithering” moves would be most efficient to resolve small registry errors, whereas inchworm moves would dominate when larger registry shifts are necessary (the exponent in Eq. 12 suggests super-diffusive scaling for long distances). However, Fig. 7 shows that our inchworm calculation predicts a success rate nearing unity for displacements less than 10 bases. Therefore, the distinction between these two kinds of moves is not significant in the regime where slithering is expected to dominate.

## Conclusion

While our model neglects many of the complexities of DNA hybridization, it provides valuable insight into key processes that have been previously reported. For instance, the slow nucleation events noted by Wetmur and coworkers^32^ are explained by the factor *P*_diff_ *P*_reg_*P*_zip_ appearing in the numerator of Eq. 3. This factor shows that, before molecules can enter the zipping phase, they must collide in an alignment compatible with an encounter complex then find the native registry. The factor *P*_zip_ will also play an important role on larger DNA molecules, which will contain many regions of intramolecular hybridization. However, once the zipping process proceeds past the first stem-loop structure, it is likely that the hybridized region will be large enough that *P*_zip_ ≃ 1. Therefore, nucleation depends on the combination of three factors: 1) a collision between unstructured regions with enough complementarity to stabilize an encounter complex, 2) the base pairs stabilizing the encounter complex must be close enough to the in-register alignment that native contacts can form within the Zimm time, and 3) the first in-register contacts must provide a strong enough anchor to stabilize the complex long enough for hybridization to work through nearby regions of non-native structure. The requirement of a strong anchor point also provides insight to the “rule of 7” observation that seven continuous base pairs provide a substantial boost to hybridization kinetics^41^.

In conclusion, we have presented a model that reduces the complex dynamics of hybridization to three basic processes. We hope that an understanding of how sequence and non-native interactions affect these events provides intuition for the design of DNA sequences for diagnostics, biotechnology, and DNA-based nanostructures.

## APPENDIX

### Zipping probability calculation

Upon the formation of the first in-register base pair, the intermolecular helix will rapidly grow to encompass the adjacent unstructured region. This means that zipping is insensitive to the precise location of the first contact, which allows us to approximate it as a zipper starting from the left end and proceeding to the right.

### Zipping probability of unstructured sequences

The probability of successfully zipping after forming *x* base pairs satisfies the convection-diffusion equation (Eq. 14). The solution to this equation is

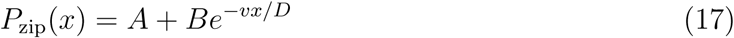

where *A* and *B* are constants. The formation of intermolecular base pairs in unstructured sequences is always favorable due to the absence of existing base pairs that need to be broken. Therefore, Δ*G* < 0 and *v* is always positive. The boundary conditions (Fig. 8) indicate failed zipping at *x* = 0 and success at *x* = *L*

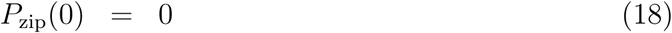

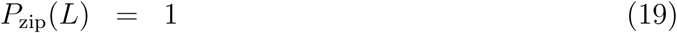

Using the boundary conditions to solve for *A* and *B* we find

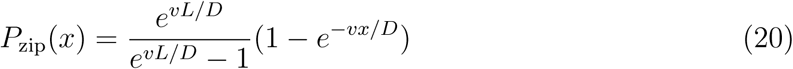

The quantity needed in Eqs. 2 and 3 is the zipping probability after the formation of the first base pair

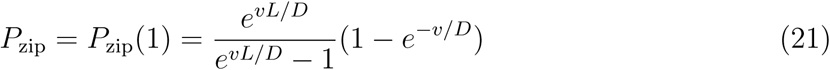

where *v* and *D* are determined from Eqs. 4, 15, and 16. The bond affinity, Δ*G*, appearing in these expression is determined by averaging the nearest-neighbor energies at 0.5M salt over the entire sequence.

### Zipping probability of stem-loop sequences

To calculate *P*_zip_ for stem-loop sequences we divide the molecule into three regions (Fig. 8). The first region, 0 < *x* < *x*_0_, is the unstructured tail which forms the first intermolecular base pairs. The second region *x*_0_ < *x* < *x*_1_ encompasses the stem-forming nucleotides adjacent to region 1. The third region, *x*_1_ < *x* < *L*, includes the loop, the stem-forming nucleotides not adjacent to region 1, and the unpaired tail (if present). Zipping in regions 1 and 3 is unimpeded by intramolecular structure, so Δ*G* is negative and *v* is positive. Zipping in region 2 requires disruption of the existing intramolecular stems, so Δ*G* is positive and *v* is negative. The solutions to Eq. 14 in each of these regions is

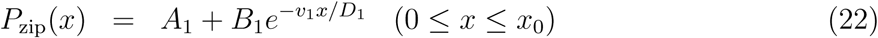

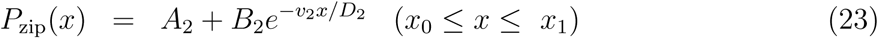

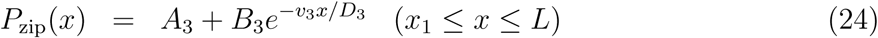

To find *A*_1_, *B*_1_, *A*_2_, *B*_2_, *A*_3_ and *B*_3_ we use the boundary conditions shown in Fig. 8.

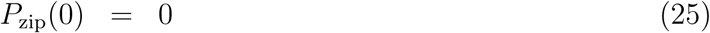

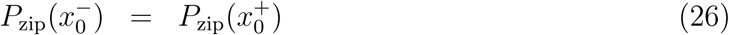

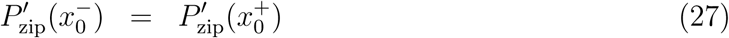

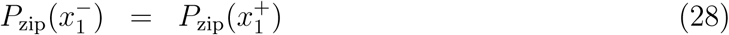

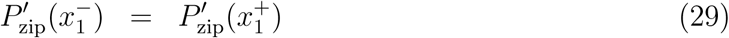

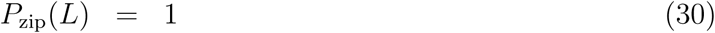

The final result is not particularly illuminating, and too unwieldy to show here, but is plotted in Fig. 8c. The drift velocities *v*_*i*_ and diffusion coefficients *D*_*i*_ are determined for each region by averaging the nearest-neighbor stabilities at 0.5M salt within that region.

### Gillespie simulation

The binding lifetimes *t*_reg_(*R*) depend sensitively on the sequence of bases at the point of first intermolecular contact. Therefore, it is necessary to account for the helix extending in both directions from the point of contact. The bidirectional extension of a helix means that a given base pair has a different Δ*G* of formation depending on whether it is on the left or right end of the growing helix because it has a different nearest-neighbor in each case. Due to this complication, we do not attempt to solve the first passage time analog of Eq. 13^45,51^, and instead, compute the binding lifetimes by Gillespie simulation^45,48,66^.

At each step of the Gillespie algorithm, the allowed moves are the formation and breakage of base pairs at either end of the helix, as shown in Fig. 13. The rate constants for each move are determined from Eq. 4 with Δ*G* computed from the nearest-neighbor model^5,17–22^.

**Figure 13:**
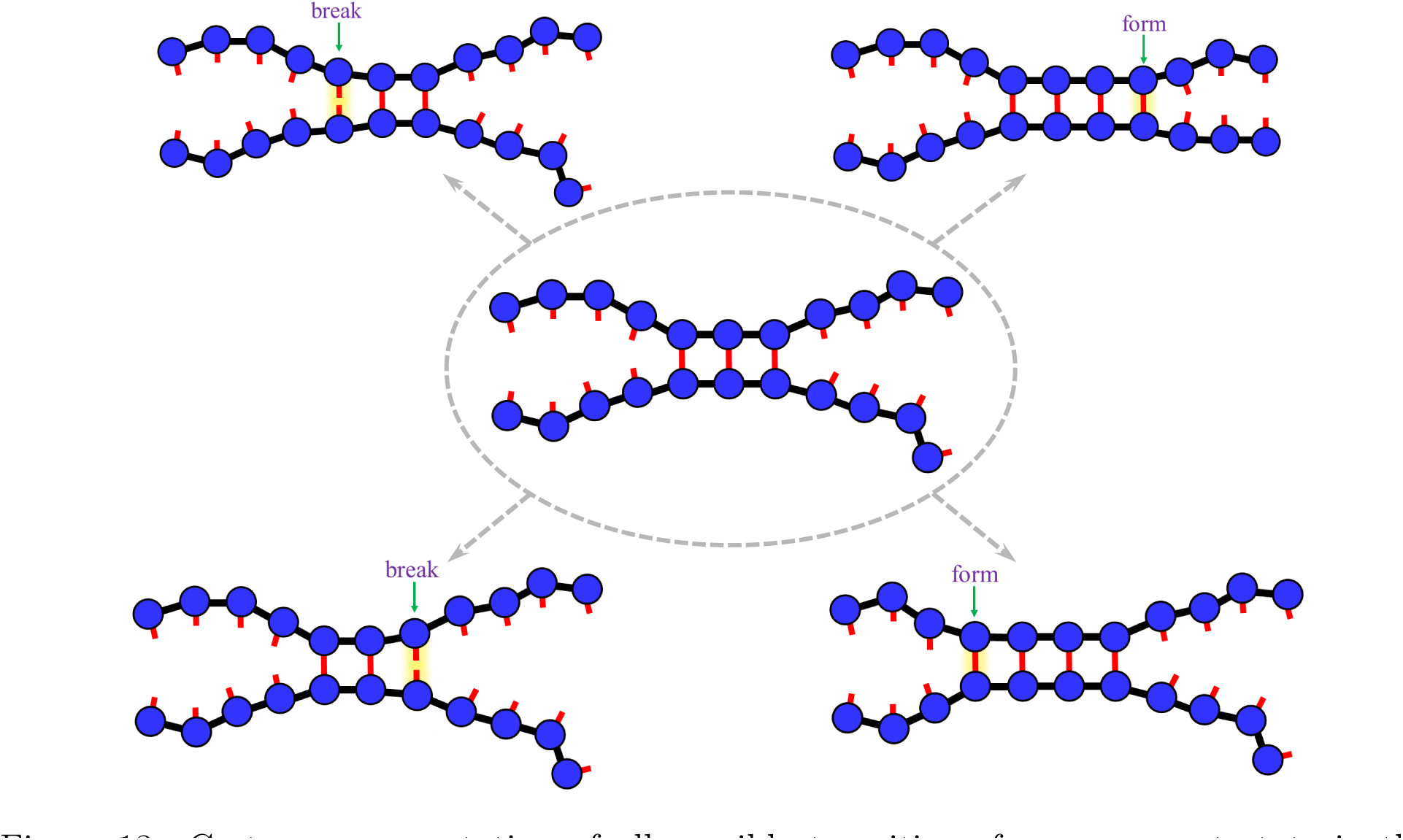
Cartoon representation of all possible transitions from a current state in the Gillespie simulation. The DNA strands can form more or break one base pair at either end of the existing base pairs.

### List of single DNA strands

The following tables show the sequences from Zhang et al.^44^ used in the development of our model. These sequences are filtered so that they contain only unstructured molecules or simple stem-loop structures in the monomer state (as predicted by NUPACK^23^). Furthermore, we only consider sequences where both strands have identical predicted structures in the monomer state.

**Table 2:**
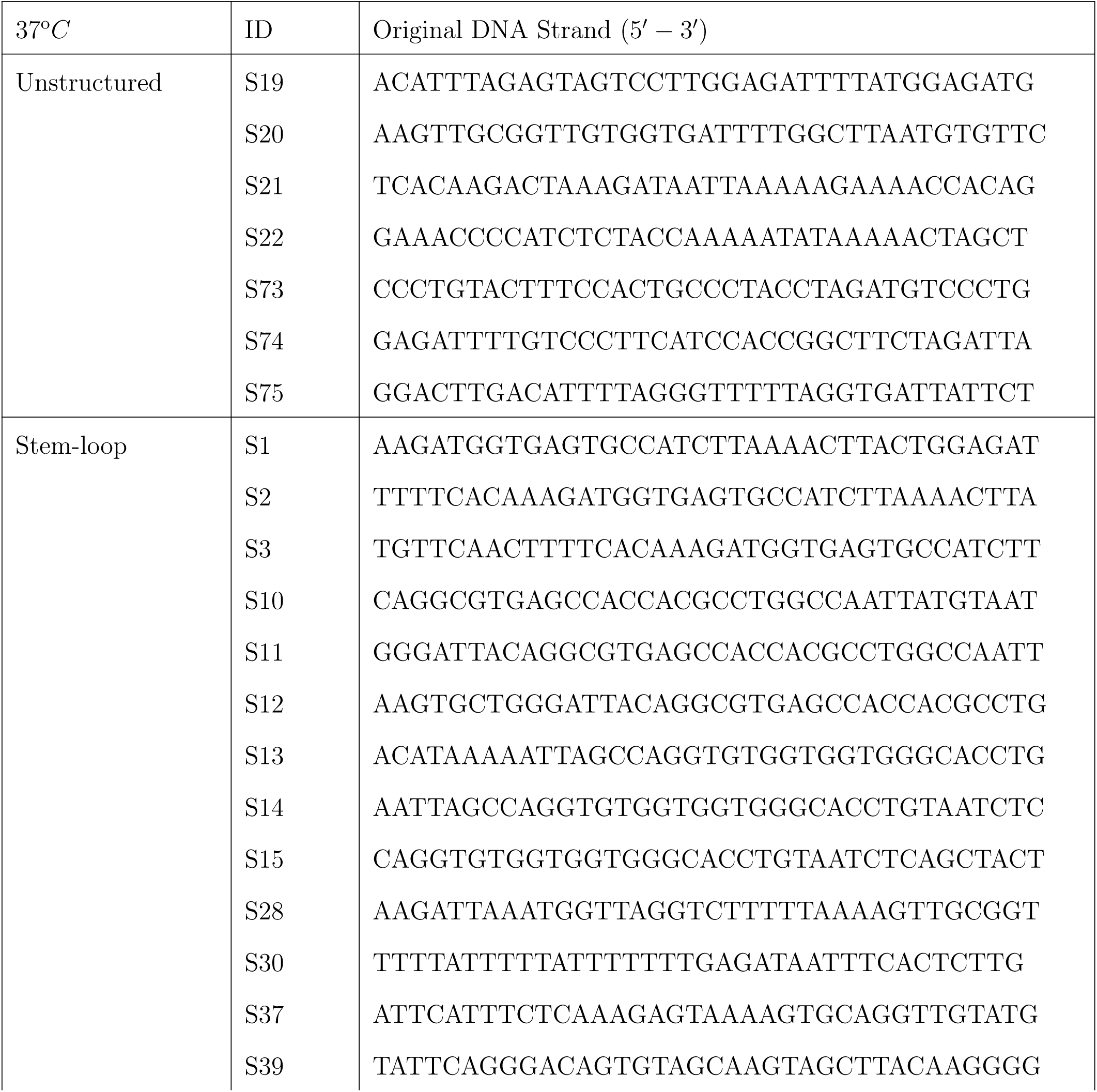
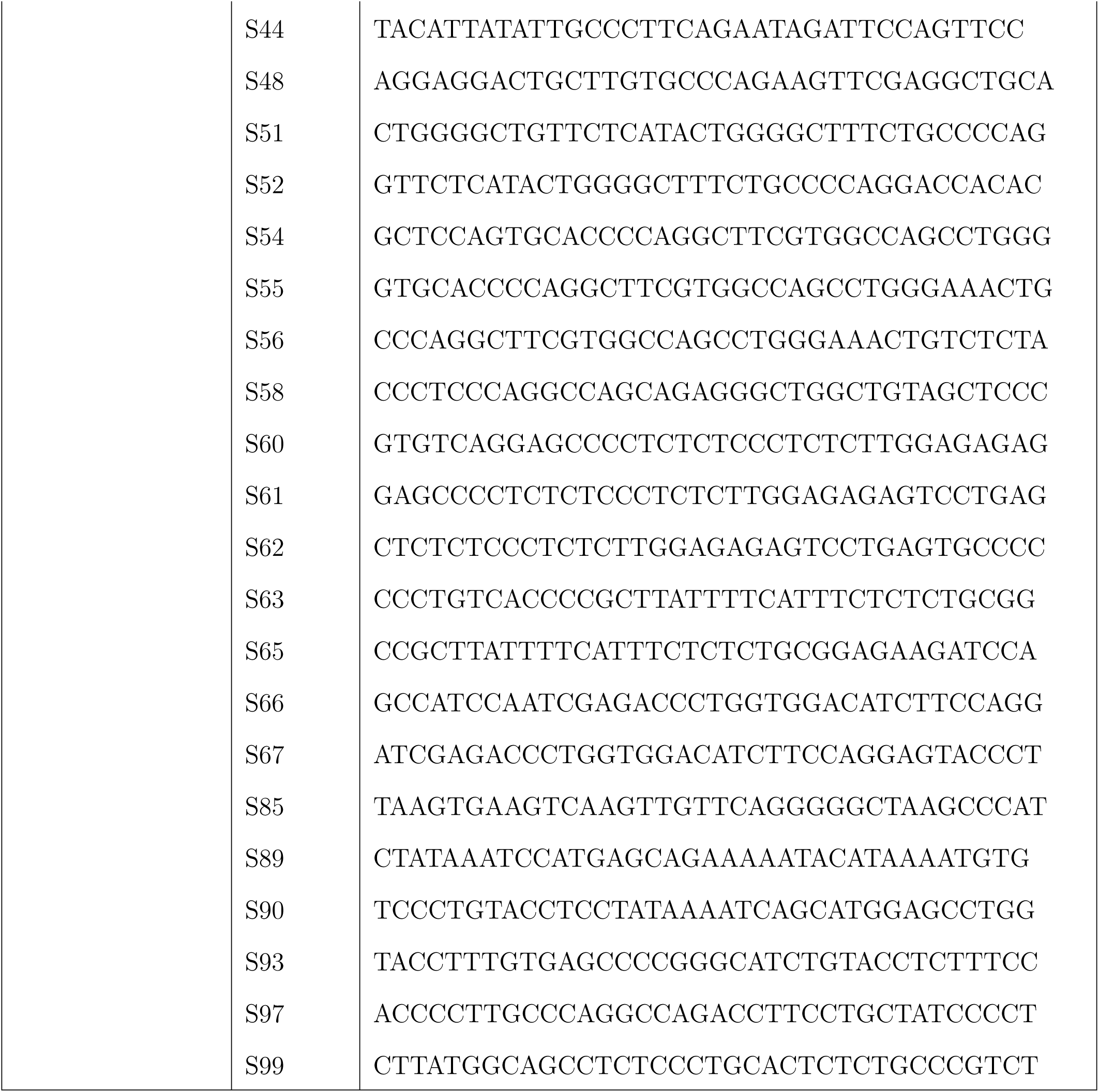
List of single DNA strands at 37°C

**Table 3:**
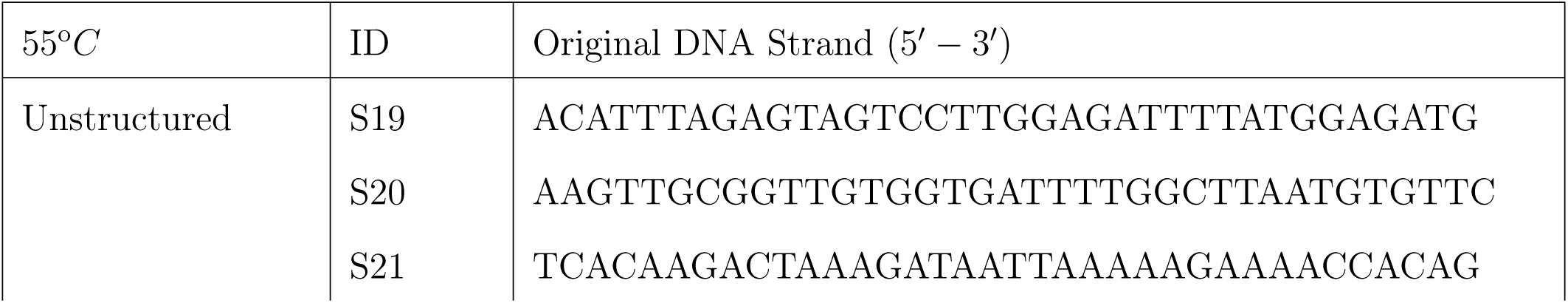
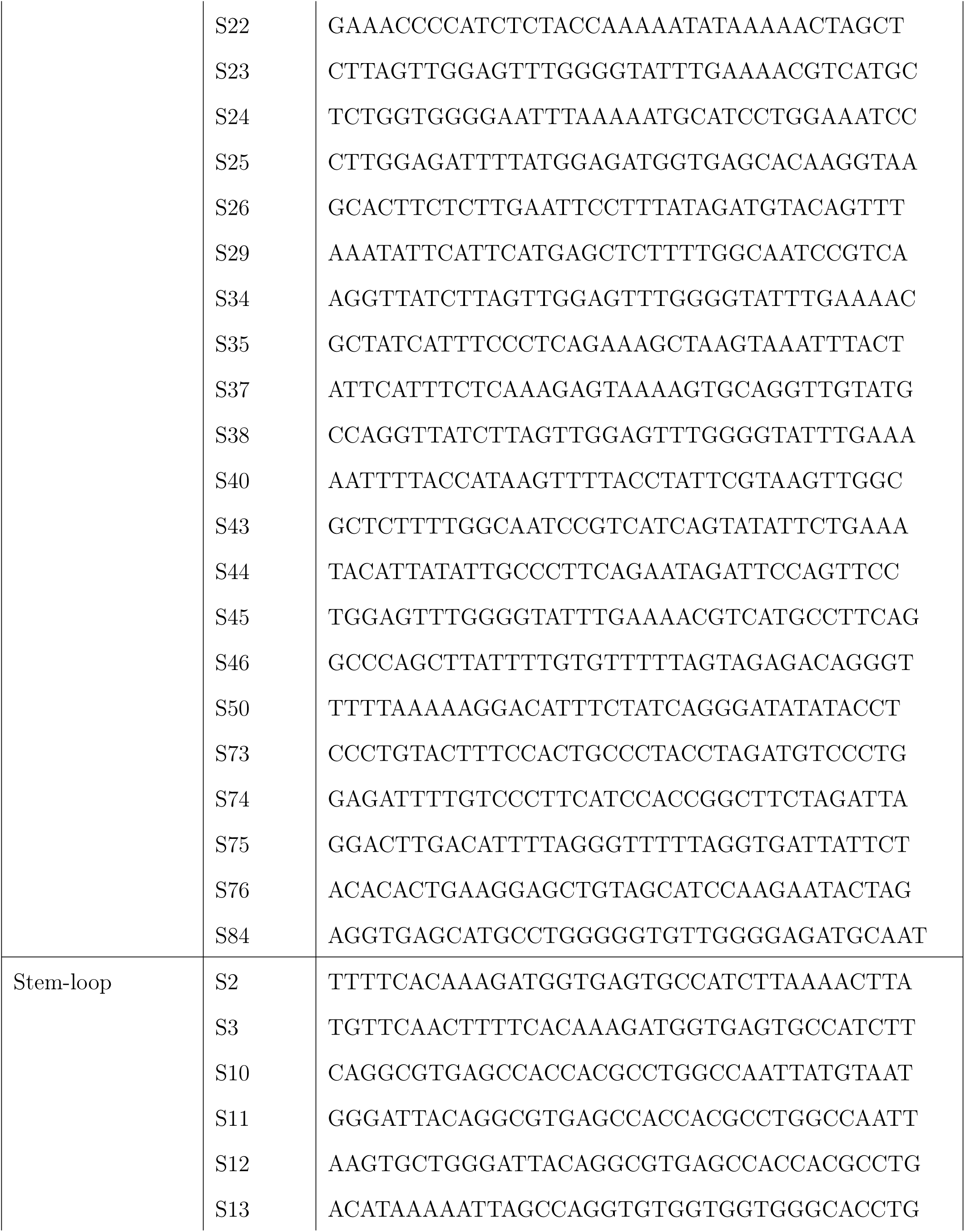
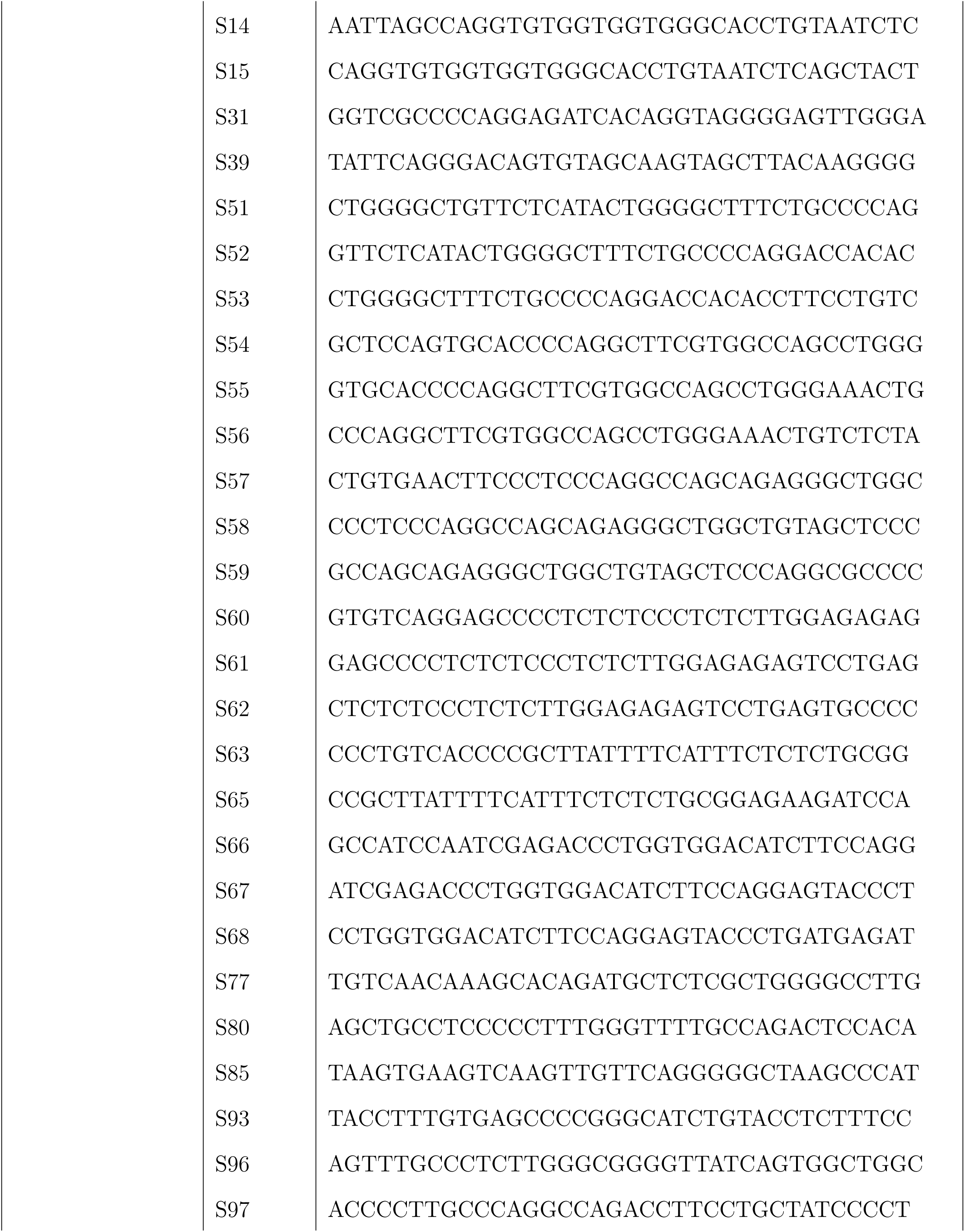
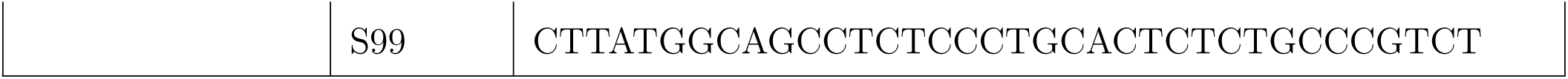
List of single DNA strands at 55°*C*

## Acknowledgement

This work was supported by National Institutes of Health grants R01GM107487 and R01GM141235.

